# Field-based prediction of sugarcane photosynthesis through environmental inputs

**DOI:** 10.1101/2025.03.11.642581

**Authors:** Rafael L. Almeida, Tamires S. Martins, José R. Magalhães Filho, Regina C. M. Pires, Marcos G. A. Landell, Mauro A. Xavier, Eduardo C. Machado, Rafael V. Ribeiro

## Abstract

Sugarcane (*Saccharum officinarum*) is a highly productive C_4_ crop prevalent in tropical and subtropical areas. However, its photosynthetic efficiency is influenced by environmental factors such as light, moisture and temperature. Understanding these interactions is critical for optimizing yields and addressing climate-related challenges. This study investigated the effects of environmental variables on carbon assimilation in four Brazilian sugarcane varieties (SP79-1011, IAC94-2094, IACSP94-2101 and IACSP95-5000), addressing both optimal and limiting conditions for key parameters. Over a 530-day field experiment, data were collected every 30 days from 7:00 to 17:00, measuring diurnal CO_2_ assimilation (*A*), photosynthetically active radiation (PAR), vapor pressure deficit (VPD), and air temperature. Polynomial models and multiple linear regression were used to quantify the contributions of these variables in CO_2_ uptake, yielding robust model fits (*p*<0.05, R^2^ = 0.84–0.99). Herein, optimal photosynthetic performance occurred under PAR at 1800 μmol m^−2^ s^−1^, VPD at 2.34 kPa, and air temperature close to 32.5°C. A strong correlation (r = 0.92, p<0.001) between observed and predicted photosynthesis and high model efficacy (R^2^=0.60, p<0.001) underscored the reliability of the approach, explaining 60% of the observed variation. While the results highlighted the model’s effectiveness in predicting sugarcane photosynthetic rates under varying diurnal and seasonal conditions, deviations indicated the influence of unmeasured parameters and complex interactions that need further investigation. These findings provide valuable insights to refine sugarcane management practices, enhance yield potential, and improve crop resilience under climate change scenarios.

## Introduction

Sugarcane (*Saccharum officinarum*) is a highly productive C_4_ crop prevalent in tropical and subtropical areas. Its C_4_ photosynthetic pathway allows efficient performance in high-light and high-temperature environments by minimizing photorespiration, giving it an advantage over C_3_ crops under such conditions. Despite this adaptation, sugarcane’s photosynthetic performance remains influenced by environmental factors, including light (Almeida, et al. 2022), moisture and temperature (Cerqueira et al. 2019; Cruz et al, 2021; Martins et al, 2024). Understanding the impact of these factors is essential for optimizing sugarcane yields and mitigating the impacts of climate-related variability on production. This study investigated the effects of environmental and physiological variables on carbon assimilation in sugarcane, addressing both optimal and limiting conditions for key parameters.

## Material and methods

### Experimental conditions and evaluations

An irrigated field experiment was conducted using four Brazilian sugarcane varieties (SP79-1011, IAC94-2094, IACSP94-2101 and IACSP95-5000), grown in a

Eutrophic Red Latosol in Campinas SP, Brazil (22°52’S, 47°04’W, 665 m a.s.l.). Over a 530-day period, data were collected every 30 days, from 7:00 to 17:00, encompassing diurnal measurements of CO_2_ assimilation (*A*) photosynthetically active radiation (PAR), vapor pressure deficit (VPD) and air temperature (T_air_). These parameters were measured using an infrared gas analyser (Li-6400XT, LICOR Inc., USA) on the first fully expanded leaf with a visible ligule (leaf +1). Air temperature (model HMP45C, Campbell, North Logan UT, USA) and rainfall (model CS700, Campbell, North Logan UT, USA) were recorded using a meteorological station installed within the experimental area. Data were logged using a CR1000 data logger (Campbell, North Logan UT, USA). The climatological water balance was computed in accordance with Rolim et al. (1998), utilizing a soil water capacity of 98 mm, as defined by Cruz et al. (2021).

### Predicting photosynthesis

To explore the relationships between leaf CO_2_ assimilation (*A*) and environmental variables such as PAR, VPD and T_air_, envelope graphs were generated by fitting quadratic or cubic polynomial functions to the upper limits of the data distribution (Fig.2). Model selection criteria included the coefficient of determination (R^2^ = 0.84−0.99), residual analysis and statistical significance of predictors (*p*<0.05). Additionally, a multiple linear regression analysis was applied to quantify the individual contributions of the predictors to *A* (model calibration; *p*<0.001). The relationship between the observed and predicted values of *A* was validated using an independent dataset, assessed through Pearson’s correlation coefficient and linear regression analysis (*p*<0.001). To ensure the robustness of the analysis, all variables were collected simultaneously and consolidated into a unified dataset. Cases with missing values were excluded to maintain the integrity of the dataset. Statistical analyses were performed using R software (R Core Team, 2024; version 4.4.0) employing the packages: ‘broom’, ‘car’, ‘dplyr’, ‘ggplot2’, ‘gridExtra’, ‘openxlsx’, ‘purr’, ‘readxl’ and ‘writexl’.

## Results and Discussion

During the experimental period, total rainfall combined with irrigation reached 2,135 mm. Air temperatures ranged from 9.3°C to 33.1°C, averaging 21°C (Fig. 1a). The plants experienced water stress during two critical phases: early in the growth cycle, when irrigation was not operational, and later during the maturation phase, when irrigation was intentionally suspended to promote sugarcane maturation.

**Fig. 1.**
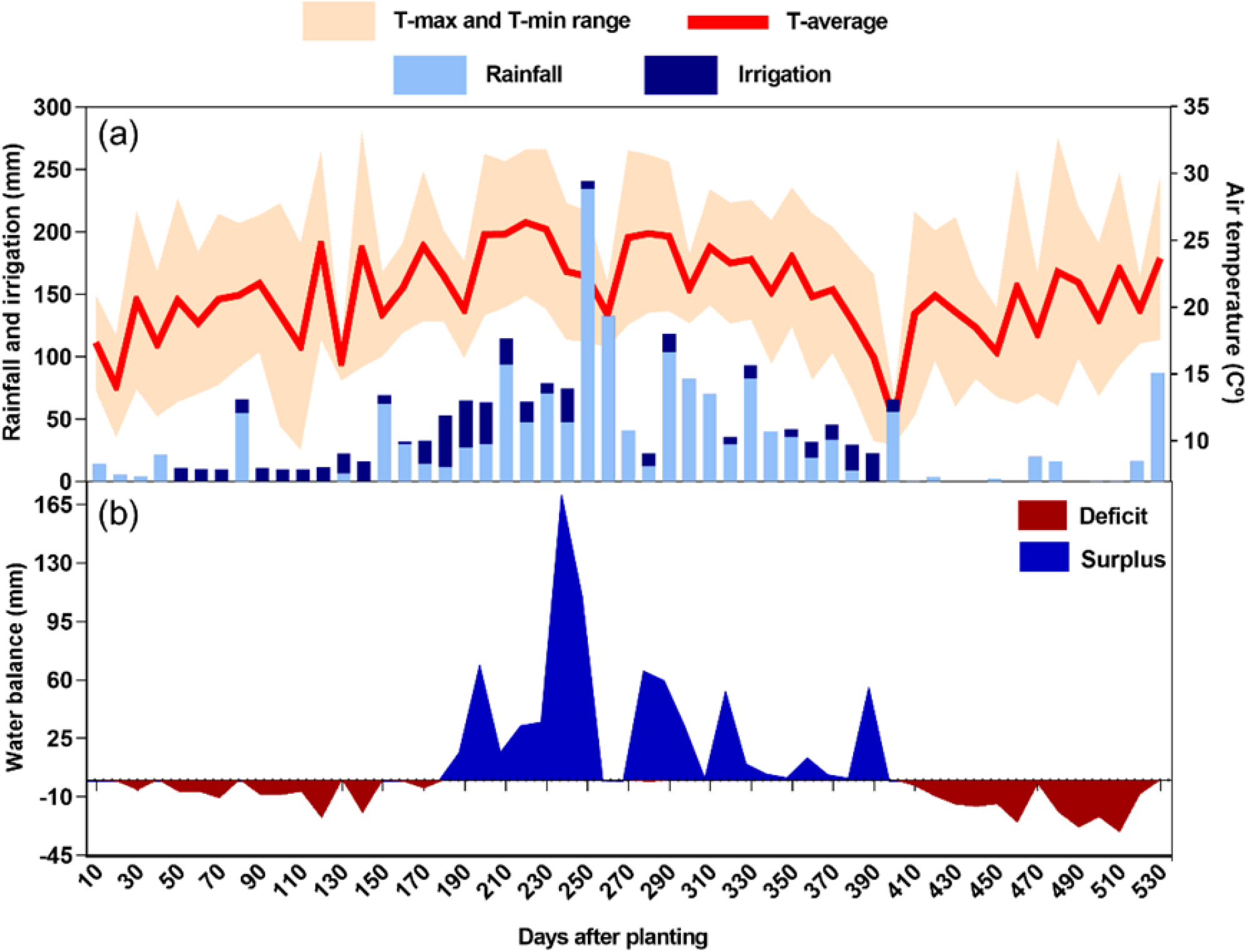
Rainfall, irrigation, and average maximum and minimum air temperatures (a), and climatological water balance (b) throughout the sugarcane growth cycle.

Photosynthesis exhibited distinct responses to environmental and physiological factors, with maximum rates observed under specific conditions. Herein, we found key environmental factors driving sugarcane photosynthesis, including PAR, VPD and T_air_. Leaf CO_2_ assimilation (*A*) increased with rising PAR, peaking at 43 μmol m^−2^ s^−1^ when PAR reached approximately 1800 μmol m^−2^ s^−1^ before plateauing and subsequently decreasing (Fig. 2a). In fact, C_4_ photosynthesis is strongly correlated with light (Zhu et al. 2010; Almeida et al. 2022). In contrast, VPD showed a different pattern: *A* peaked at 42 μmol m^−2^ s^−1^ around 2.34 kPa, followed by a sharp decline as VPD increased further (Fig. 2b). This suggests that excessive atmospheric moisture demand is compromising carbon assimilation. Like VPD, T_air_ exhibited bell-shaped response, with *A* reaching a maximum of 44.5 μmol m^−2^ s^−1^ at an optimal temperature of 32.5°C (Figs. 2c). These results highlight the complex interplay between environmental and physiological factors, demonstrating that sugarcane photosynthesis operates most efficiently within specific ranges of PAR, VPD, and temperature. Deviations from these optimal conditions reduce the photosynthetic performance, emphasizing the importance of maintaining favorable growth conditions to maximize carbon assimilation. Finally, the strong alignment of observed and predicted *A* data points along the 1:1 line of the model in Fig. 3, demonstrates its predictive accuracy, with a higher correlation (r=0.92, *p*<0.001). Herein, the model explained 60% of the variation in *A*, attributable to PAR, VPD and T_air_.

**Fig. 2.**
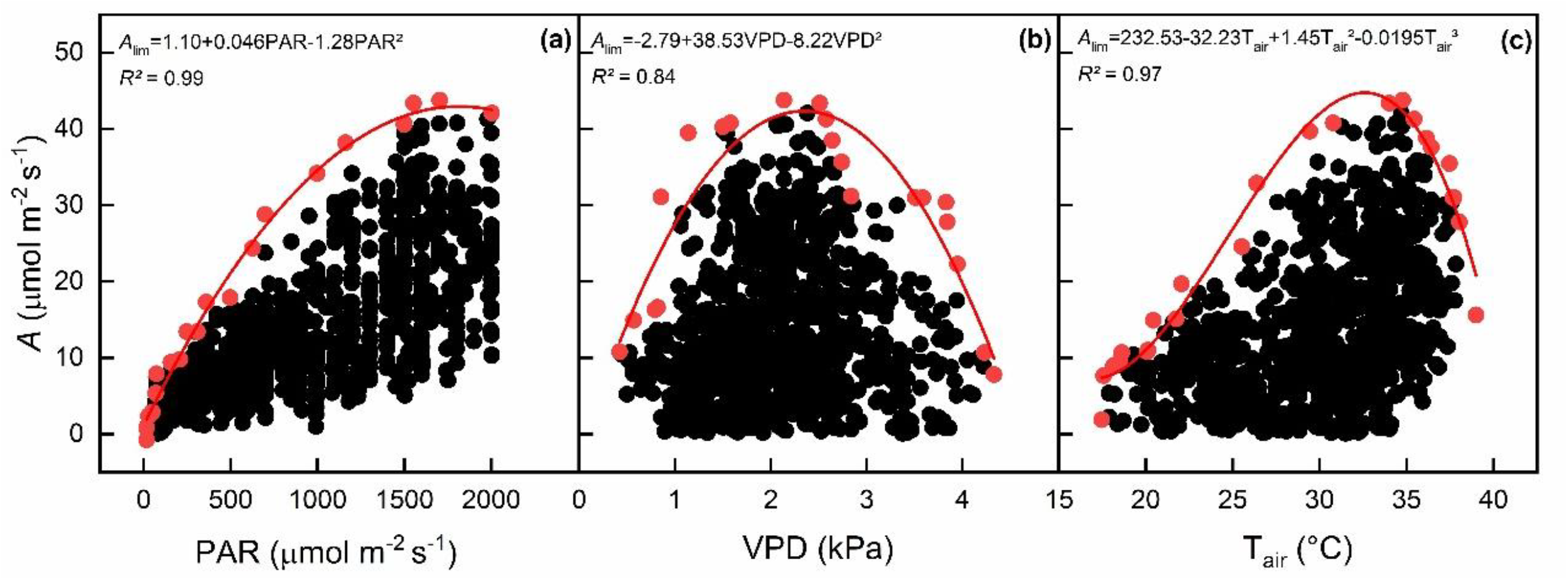
Relationship nbetween photosynthetic rate (*A*) and photosynthetic active radiation (PAR, in a), vapor pressure deficit (VPD, in b) and air temperature (T_air_, in c) throughout the sugarcane growth cycle. The red line represents the fitted model (quadratic or cubic), while the red dots indicate the data points used for fitting. R^2^ is the adjusted coefficient of determination.

**Fig. 3.**
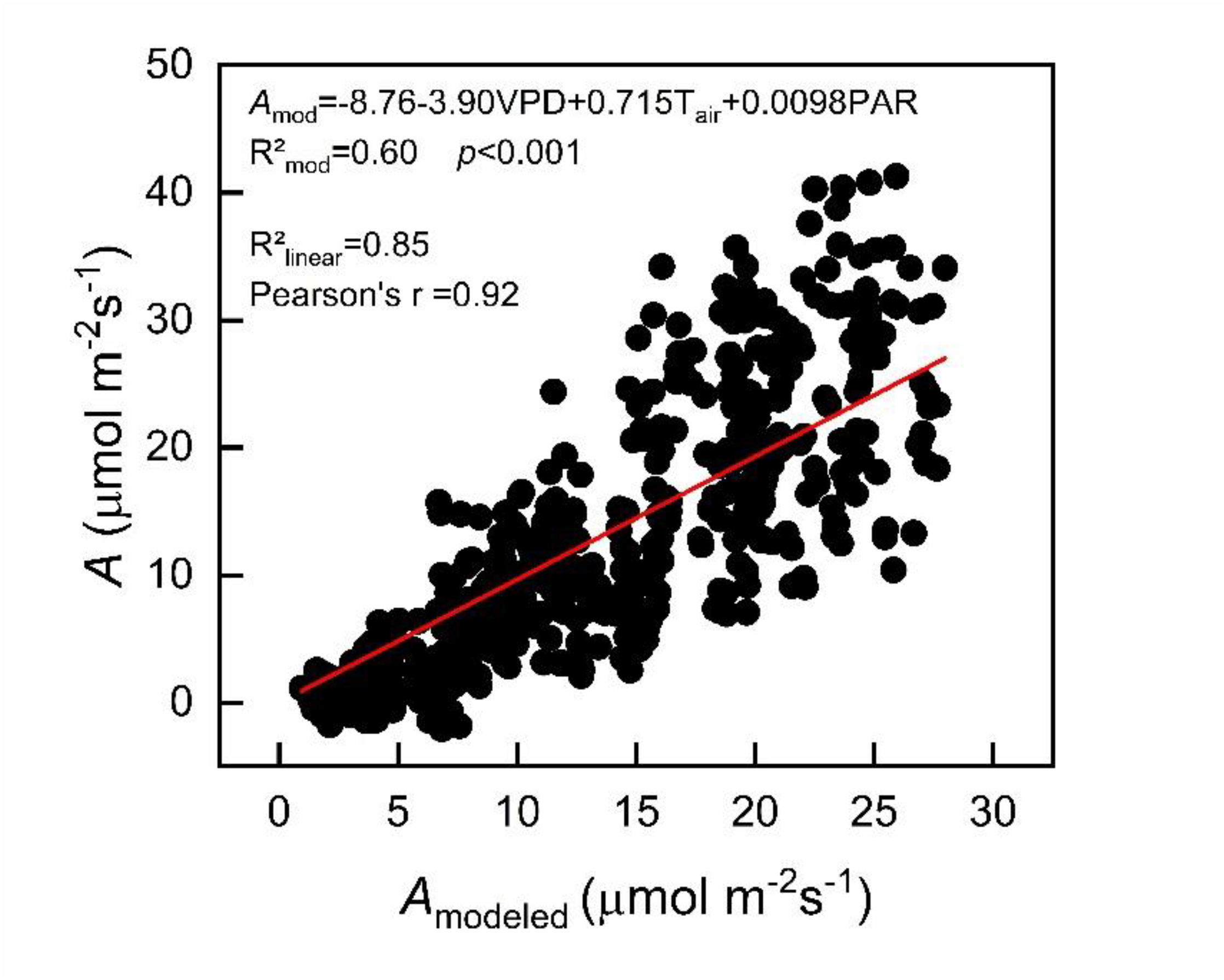
Relationship between photosynthetic rate (*A*) and modeled photosynthetic rate (*A*_modeled_) throughout the sugarcane growth cycle. The red line represents the fitted linear regression. R^2^ is the adjusted coefficient of determination and r is the Pearson’s correlation coefficient.

## Conclusion

Our findings demonstrate the strong predictive capacity of the proposed model in estimating sugarcane photosynthetic rates across varying diurnal and seasonal environmental conditions. Despite its robust performance, observed deviations highlight the need for further investigation to address potential limitations, such as unmeasured variables or intricate interactions affecting photosynthesis. Understanding the interactions provides valuable insights for refining sugarcane management strategies. By integrating these variables into predictive models like the one presented here, we can optimize plant growth, enhance yields, and strengthen the resilience of sugarcane cultivation in the face of evolving climate change scenarios.

